# Can citizen science analysis of camera trap data be used to study reproduction? Lessons from Snapshot Serengeti program

**DOI:** 10.1101/2020.11.30.400804

**Authors:** Thel Lucie, Chamaillé-Jammes Simon, Keurinck Léa, Catala Maxime, Packer Craig, Sarah E. Huebner, Bonenfant Christophe

## Abstract

Ecologists increasingly rely on camera-trap data to estimate biological parameters such as population abundance. Because of the huge amount of data, the assistance of non-scientists is often sought after, but an assessment of the data quality is necessary. We tested whether volunteers data from one of the largest citizen science projects - Snapshot Serengeti - could be used to study breeding phenology. We tested whether the presence of juveniles (less than one or 12 months old) of species in the Serengeti: topi, kongoni, Grant’s gazelle, could be reliably detected by the “naive” volunteers vs. trained observers. We expected a positive correlation between the proportion of volunteers identifying juveniles and their effective presence within photographs, assessed by the trained observers.

The agreement between the trained observers was good (Fleiss’ κ > 0.61 for juveniles of less than one and 12 month(s) old), suggesting that morphological criteria can be used to determine age. The relationship between the proportion of volunteers detecting juveniles less than a month old and their actual presence plateaued at 0.45 for Grant’s gazelle, reached 0.70 for topi and 0.56 for kongoni. The same relationships were much stronger for juveniles younger than 12 months, reaching 1 for topi and kongoni. The absence of individuals < one month and the presence of juveniles < 12 months could be reliably assumed, respectively, when no volunteer and when all volunteers reported a presence of a young. In contrast, the presence of very young individuals and the absence of juveniles appeared more difficult to ascertain from volunteers’ classification, given how the classification task was presented to them.

Volunteers’ classification allows a moderately accurate but quick sorting of photograph with/without juveniles. We discuss the limitations of using citizen science camera-traps data to study breeding phenology, and the options to improve the detection of juveniles.

## Introduction

Camera trapping is increasingly used for ecological monitoring due to its low cost, relative ease of use, and the variety of data it can supply (O’Connell et al. 2010). For instance, camera trap data are used to study species’ occupancy and co-occurrence (Anderson et al. 2016), population dynamics (Karanth et al. 2006), or individual behaviour (*e.g.* vigilance behaviour: Chamaillé-Jammes et al. 2014, or diel activity patterns: Luo et al. 2019). A potential drawback of camera traps is the huge amounts of data that are generated (> 7 millions photographs for the Snapshot Serengeti initiative alone). Ecologists have realized that the benefits of continuously collecting data in the field can quickly be negated by the burden of database management, and visual inspection and analysis of photographs to record the desired data (Wearn and Glover-Kapfer 2017).

To process such a massive amount of information in a reasonable time, scientists have sought the help of non-specialists who perform diverse tasks like counting objects in photographs, describing picture content or identifying animal and plant species (*e.g.* McShea et al. 2015). Initially, part of the scientific community was sceptical about citizen science, in particular questioning data quality (Riesch and Potter 2014). However, volunteers have sometimes proven to be as efficient as experts, as for instance for the identification of large herbivore species in savanna ecosystems (Swanson et al. 2016). More recently, the advances in deep learning have led computers to become as efficient as people at identifying species, and, sometimes, behaviour classification problems (Norouzzadeh et al. 2018). However, human judgment is still valuable in particular cases where too little data are available to train models (*e.g.* active learning, Joshi et al. 2009), or when differences among the objects to be classified are subtle and classification requires some subjectivity (Miele et al. 2020, *in prep.*). We believe this is the case for age classification problems, for which to the best of our knowledge the number of precisely labelled pictures is currently too small to allow an efficient and reliable automatization of the process.

Under the assumption that the detectability of juveniles and adult females segments of the population is not biased by camera traps methodology, classifying individuals into ageclasses such as juveniles and adults would allow estimates of key demographic parameters (*e.g.* reproductive rates) or life-history traits (*e.g.* breeding phenology), which are essential in estimating population growth rates (Sibly and Hone 2002). For instance, Ogutu and colleagues (2008) highlighted that rainfall influences the abundance of several large herbivore species of the Mara-Serengeti ecosystem, by acting differently on each segment of the populations at specific periods of the year. Furthermore, it would facilitate the study of the relationships between population characteristics and their environments such as between birth phenology, diet and food resource availability (Sinclair et al. 2000), or their potential evolution in the context of climate change (Visser and Both 2005). Until now, the study of those key demographic parameters has been mainly conducted by direct field observations (*e.g.* Côté and Festa-Bianchet 2001 in mountain goats, Plard et al. 2013 in roe deer). However, this methodology still requires an intensive and often costly field effort. Identifying and counting juveniles from camera traps could reduce this field effort, or allow larger-scale or longer-term studies, as suggested by Hofmeester et al. 2019, but could also be timeconsuming because of tedious data processing. With the help of citizen science, data handling time could be substantially reduced, but the accuracy of non-specialists in detecting juveniles of large mammals from photographs has not yet been explored.

Here, we evaluate the usefulness of camera trap data annotated by citizen scientists online to assess the presence of juveniles of large herbivores in the photographs. We use photographs and citizen classifications from the Snapshot Serengeti project (Swanson et al. 2015), one of the world’s largest citizen science programs, on a subset of the data. We focus on the detection of juveniles in three species found in the Serengeti National Park, Tanzania with contrasting social and neonatal behaviours: topi (*Damaliscus jimela*), kongoni (*Alcelaphus cokii*) and Grant’s gazelle (*Nanger granti*). We first evaluate the agreement between trained observers from our research team, and then test the ability of the volunteers to detect juveniles by comparing their classification with ours. We predict a better agreement between trained observers for the youngest age class because determination criteria are clearer and easier to identify than for older juveniles (*e.g.* absence of horns). Consequently, the level of agreement should decrease for age classes that are based on more subjective or difficult-to-assess criteria (*e.g.* shape of the horns). Under the hypothesis that volunteers could generally identify juveniles correctly, we expect a positive relationship between the proportion of volunteers reporting a juvenile on a photograph and the probability of the actual presence of a juvenile, as determined by the trained observers. Again, we expect the correlation to be stronger for the youngest age class of juveniles because they are easier to differentiate from adults. Across species, we expect a higher agreement and correlation for topi and kongoni than for Grant’s gazelle because the former are larger, live in smaller groups and have similar body growth rate between males and females (Wilson and Mittermeier 2011), hence reducing the risk of confusion between young males and older females. Overall, our study details the strengths and weaknesses of camera trap data, in particular when classified by citizen scientists, for the study of reproductive traits such as reproductive rates or breeding phenology.

## Materials and methods

### Study site

The surveyed area within the Serengeti National Park, Tanzania, is composed of open plains and savanna woodlands. Rain mostly occurs between November and June (wet season), with mean annual rainfall increasing from 500 mm in the southeast to 1,100 mm in the northwest. This area harbours a rich community of large herbivores, composed of gregarious and migratory wildebeest (*Connochaetes mearnsi*), zebra (*Equus sp.*) and Thomson’s gazelle (*Eudorcas nasalis*), but also resident populations such as Cape buffalo (*Syncerus caffer*) or warthog (*Phacochoerus africanus*) (Sinclair and Norton-Griffiths 1995). Community dynamics are driven both by herbivores, maintaining an open state of the grassland by intensive grazing (McNaughton 1985, Sinclair and Norton-Griffiths 1995) as well as large predators (*e.g.* lion (*Panthera leo*) and hyena (*Crocuta crocuta*), Sinclair and Norton-Griffiths 1995).

### Camera trap data

The Snapshot Serengeti camera trap grid was deployed in 2010 in Serengeti National Park, Tanzania, to monitor lions and their prey, though the bycatch of numerous other species has proven useful as well. Running continuously since 2010, the grid spans 1,125 km^2^ in the center of the park. We used data provided by Snapshot Serengeti camera survey recorded between July 2010 and April 2013 (Supporting information 1). The camera traps were set ~ 50 cm above ground in the centre of a 5 km^2^ grid cell. The detection radius was approximately 45° and their field of view about 14 m (Swanson et al. 2015). Cameras took a rapid series of three pictures upon trigger of the motion and heat sensors (“capture event” in Swanson et al. 2015, hereafter called a “sequence” following Meek et al. 2014) in a few seconds interval, with a one-minute delay between sequences.

### Choice of studied species and sorting steps of the dataset

Among the many large herbivore species present in the study site, we selected topi, kongoni and Grant’s gazelle due to their contrasting biology and characteristics useful to assess the age classes of individuals. The criteria considered were (1) number of available sequences, (2) relatively small group size, (3) presence of horns in males and females, (4) relatively large size of the young (young of larger species are larger, and therefore criteria like horns are easier to detect), (5) contrasting anti-predator of behaviour of the young (Supporting information 2).

We selected the final dataset (*n* = 2,359 sequences) to conduct the analyses following several sorting steps, based on the detection of the species of interest and of the presence of juveniles from the initial complete dataset (*n* = 1,184,657 sequences) by the volunteers. We then corrected this dataset thanks to the trained observers reclassifications (see details in Table 1).

**Table 1:**
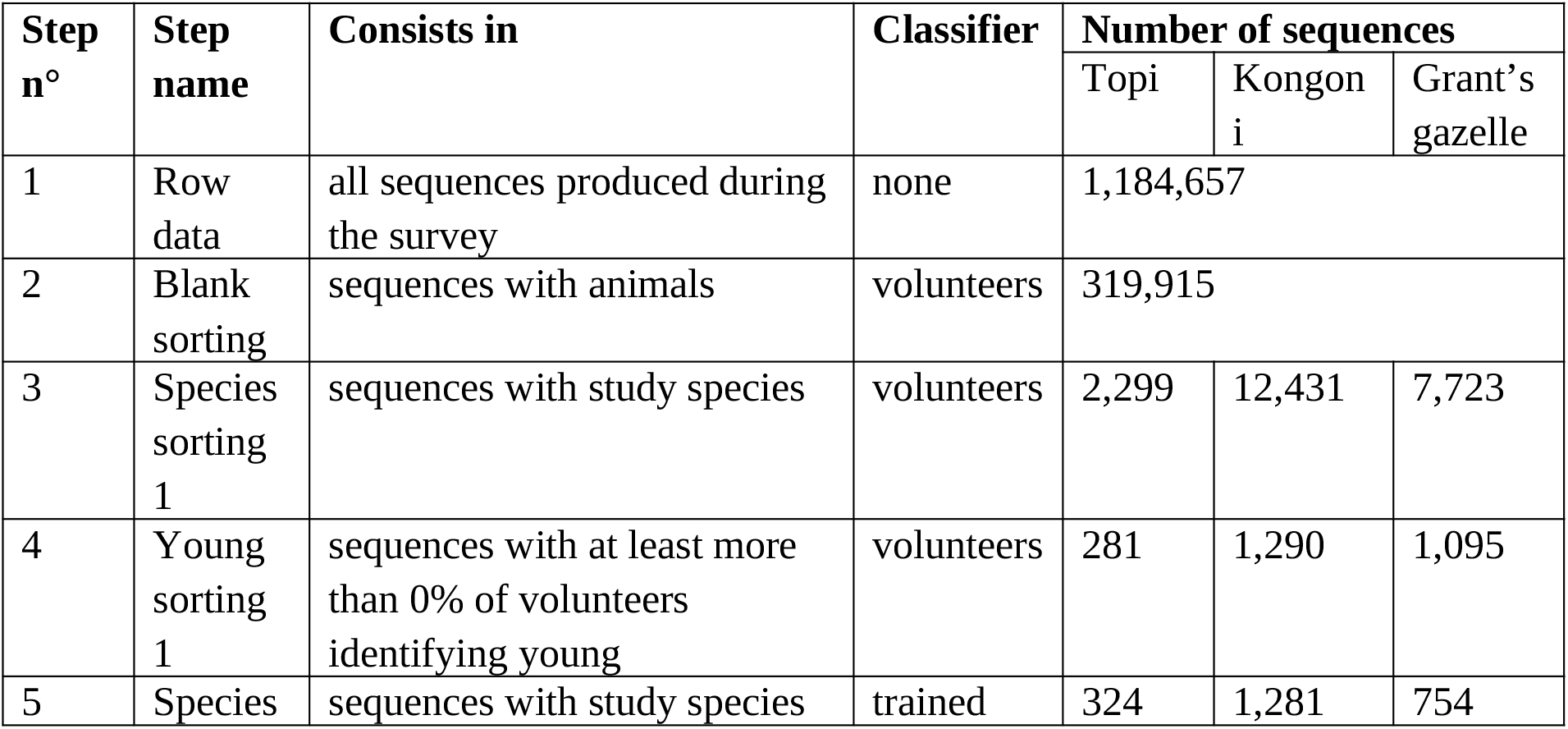
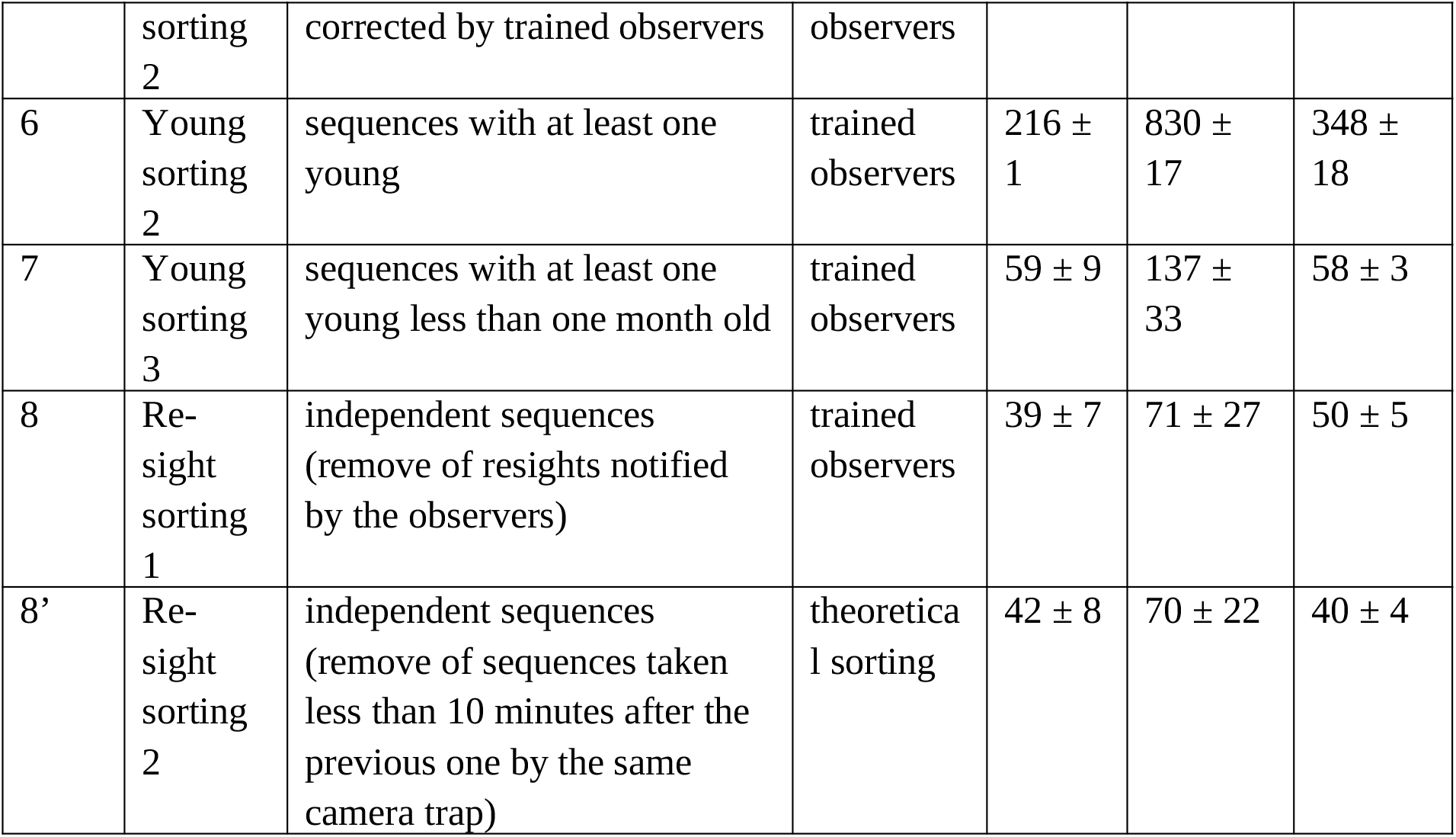
number of sequences at each sorting step from the extraction of raw data to the selection of all the independent sequences with at least one individual < 1 month old, for the three species of the study: topi, kongoni and Grant’s gazelle (pictures from Snapshot Serengeti program, Tanzania, between July 2010 and April 2013). Standard deviations are calculated on the basis of the classifications of the three trained observers.

### Assessing the presence or absence of juveniles in photographs

All Snapshot Serengeti photographs have been uploaded to the online citizen science platform “The Zooniverse” (www.zooniverse.org) to be classified by volunteers. Each sequence was classified by as many as 25 volunteers (*minimum* = 11, *maximum* = 57, Swanson et al. 2016), who each identified what species was present in the sequence and recorded whether at least one juvenile was present or not. Note that no formal definition of a juvenile was provided to volunteers, nor were there any particular guidelines about how to identify a juvenile. Volunteers simply searched and checked “young” in the Zooniverse interface when they suspected the presence of non-adult individuals. Regarding age classes, the only question volunteers had to reply to was: “Are there any young present?”. For each sequence, the volunteers’ classifications were then compiled via a plurality algorithm to yield a consensus classification, leading to a proportion of volunteers (*Pv*) who identified at least one juvenile in each sequence (see details in Swanson et al. 2015). Here, we used all sequences where volunteers identified topi, kongoni, and Grant’s gazelle, with at least one volunteer (*Pv* > 0%) having annotated the presence of a juvenile. We could not analyse all sequences for which no volunteer had reported a juvenile, as these were too many (*n* = 2,018, 11,141 and 6,628 for topi, kongoni and Grant’s gazelle respectively) to be reviewed individually. However, we checked a subset of them (*n* = 1,000 for each species), and the chance that a trained observer observed a “true” juvenile (*i.e.* of less than 12 months old, see age classes definition below) was < 6.5% for all three species studied. We did not correct observations for recaptures of the same individuals as we were only interested in the ability of volunteers to detect the presence of juveniles on the sequences, but not the actual number of juveniles.

Three of us (LT, LK and MC), considered here as trained observers, searched all sequences retained for juveniles, which were assigned to an age class when detected. We used previously published morphological descriptions of the studied species (*e.g.* shape and size of horns, size relating to the adult; Supporting information 3) to identify and age individuals. We distinguished between (1) juveniles < one month, (2) between one and six months, (3) between six and 12 months, (4) between 12 and 24 months, termed yearling hereafter and (5) individuals over 2 years old, termed adults hereafter. We defined age classes according to biological characteristics relevant to juvenile identification for each species (*e.g.* very young individuals for birth phenology identification, juveniles under one year for recruitment estimation). We recorded observers’ classifications with Aardwolf software (Krishnappa and Turner 2014). Ultimately, we produced a dataset describing the presence or absence (*M_i,s,j_* in each sequence of individuals of any of the five age categories *i*, for the species *s*, by trained observer *j*.

### Statistical analyses

We first evaluated the agreement between the three trained observers on the detection of individuals assigned to each age class for each species. We measured this agreement with the Fleiss’ κ, implemented in the *“raters”* R package (Quatto and Ripamonti 2014). Fleiss’ κ (Fleiss 1971) is the comparison of agreement between 2+ judges and the level of agreement expected by chance alone. It takes values between −1 and 1, values < 0 indicating an agreement lower to what could be expected by chance, values > 0 indicating a greater agreement than expected by chance, and values = 0 indicating an agreement close to random. We tested for significance of the difference between the Fleiss’ κ using a bootstrap procedure following Vanbelle and Albert (2008) (Supporting information 4).

We tested the relationship between the proportion of volunteers identifying at least one juvenile (*Pv*) and the probability that trained observers had identified at least one juvenile < 1 month (category *i* = 1). We also explored the same relationship with juveniles < 12 months (therefore including juveniles of categories 1 to 3 above). We fitted three generalized estimating equation models: one linear (see equation 1 below) and two piecewise models. The first piecewise model was characterized by a slope on both sides of the threshold (equation 2), the second by a slope before and a plateau after the threshold (equation 3). We fitted piecewise models to search for a potential “saturation” phenomenon, whereby beyond a specific proportion of volunteers the probability to effectively observe a juvenile does not increase anymore. We also fitted the null model for comparison. All the models were fitted for the two age classes and for each species individually, using the *wgee* function implemented in the *“wgeesel”* R package (Xu et al. 2018). We selected the best model using the Quasi-likelihood under the Independence model Criterion QIC (thresholds selected by comparison of QIC of the models for each species and age class as well). It is a modification of the Akaike Information Criterion AICc, suitable when quasi-likelihood is used instead of likelihood (Pan 2001), implemented in the *“MuMIn”* R package (Barton 2019). We used a logit link function and a binomial distribution of errors (Agresti 2002), considering the proportion of volunteers identifying at least one juvenile as fixed effect, and the identity of the sequence as clustering variable with an exchangeable correlation structure:

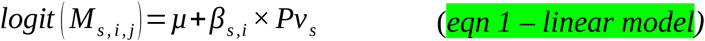

and

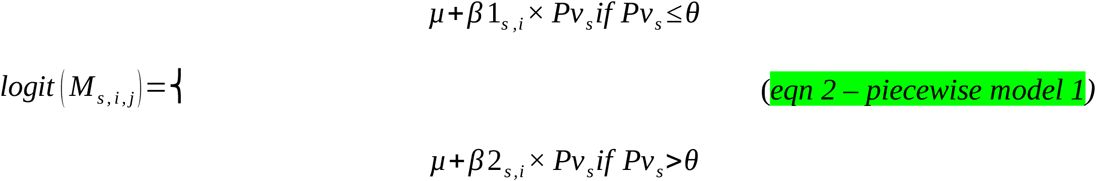

and

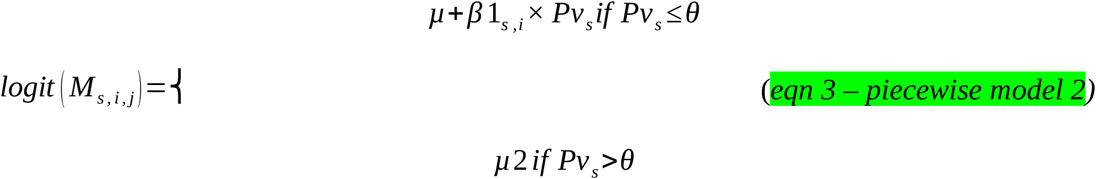

where *M_s,i,j_* denotes the presence or absence of at least one juvenile of the age class *i* (< 1 or 12 months) for a given sequence, for the species *s* and the observer *j*, *μ* is the expected mean probability of actual presence of at least one juvenile of the age class *i* when none of the volunteers identified the presence of a juvenile, *Pvs* is the proportion of volunteers identifying at least one juvenile for a given sequence, for the species *s* and *θ* is the best threshold. We performed all analyses using the R statistical software (R Core Development Team 2019).

## Results

From the Snapshot Serengeti monitoring program, we obtained 281, 1,290 and 1,095 sequences for topi, kongoni and Grant’s gazelle respectively (Fig. 1a) sorting step “Young sorting 1” and Table 1). Our selection process led to a dramatic drop in the number of usable sequences (*i.e.* those containing at least one < 1 month old juvenile), with a tenfold reduction at the species level: it only remained between 7.7% and 18.1% of the complete dataset available for the species (*n* = 59 ± 9 SD, 137 ± 33 SD and 58 ± 3 SD for each species respectively, Fig. 1a) sorting step “young sorting 3” and Table 1). The largest losses of sequences happened during the phases of selection of sequences containing the studied species and then juveniles (Fig. 1a) and Table 1). Fig. 1b) suggests that we would obtain similar results concerning the loss of sequences after selection for sequences containing juveniles for any large herbivore species recorded in the Snapshot Serengeti program.

**Figure 1:**
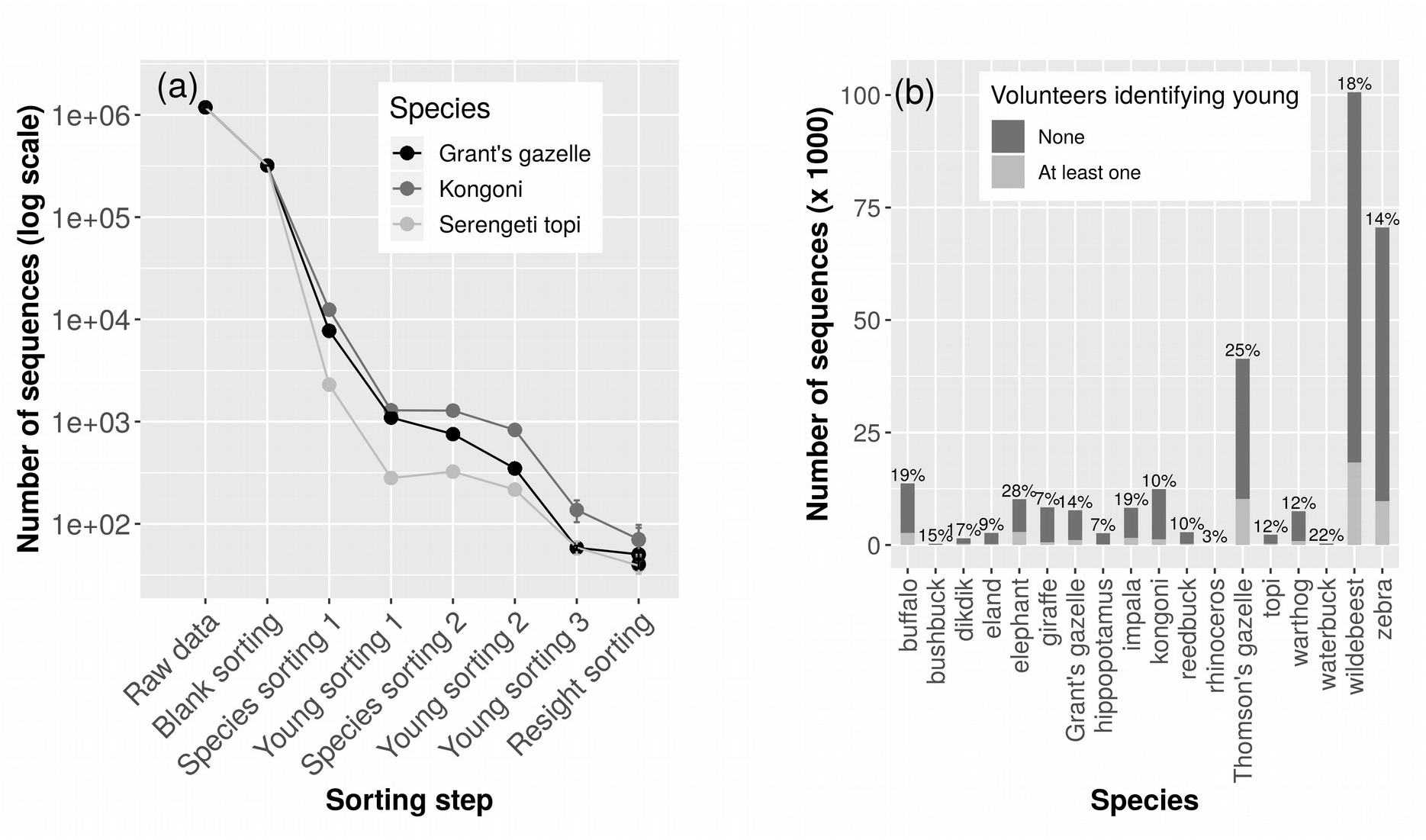
a) number of sequences available at each sorting step from the extraction of raw data to the selection of all the independent sequences with at least one individual < 1 month old, for the three species of the study: topi, kongoni, Grant’s gazelle. Pictures from Snapshot Serengeti, Tanzania, between July 2010 and April 2013. Raw data: all sequences produced during the survey; blank sorting: sequences with animals; species sorting 1: sequences with study species according to the volunteers; young sorting 1: sequences with at least one volunteer identifying “young”; species sorting 2: sequences with study species corrected by trained observers; young sorting 2: sequences with at least one young according to the trained observers; young sorting 3: sequences with at least one young < 1 month old according to the trained observers; resight sorting: independent sequences once sequences taken less than 10 minutes after the previous one by the same camera trap and presenting the same species have been removed (following Palmer et al. 2018). Note the log scale for the ordinate axis, vertical bars represent the standard deviations. b) Number of sequences available for all the large herbivore species present in the study site. The proportions indicate the proportion of sequences where at least one volunteer identified juveniles for a given species.

The three trained observers identified 993 ± 49 SD, 3,020 ± 180 SD and 2,128 ± 224 SD individuals of any age class. On average, they could not assign an age class to ~11% (n = 118 ± 68 SD), ~20% (n = 775 ± 256 SD) and 32% (n = 1,026 ± 492 SD) of the individuals for topi, kongoni and Grant’s gazelle respectively, a significant between-species difference (*χ^2^* = 262.83, *df* = 2, *p* < 0.001).

For all species, and as expected, the agreement between trained observers was highest for the youngest age class, then declined with age until the yearling class was reached (Fig. 2). Yearlings were reasonably well classified in kongoni, but overly misclassified in topi and Grant’s gazelle (Fig. 2). As expected, the agreement among trained observers was the highest for the youngest age class of juveniles, but also for the second age class and the adults, with Fleiss’ κ values almost always > 0.61 (denoting a substantial agreement, following Landis and Koch 1977), except for Grant’s gazelle aged between one and six months and adult topi. Agreement among observers was the highest for topi aged < 1 month old (Fleiss’ κ = 0.78 [0.72; 0.84]). In support of our prediction, we observed the lowest agreement for juveniles aged six months and older, and more obviously so for Grant’s gazelle. Agreement among observers concerning the three first age classes pooled together (representing the juveniles) was very good, with Fleiss’ κ values largely > 0.61 for all the species. The same holds true for juveniles between one and 12 months old when pooled together. Our results were globally consistent among the three species studied, with the highest agreement for topi, and the least for Grant’s gazelle (Fig. 2). All estimated Fleiss’ κ were significantly different from an agreement obtained by chance (Supporting information 4).

**Figure 2:**
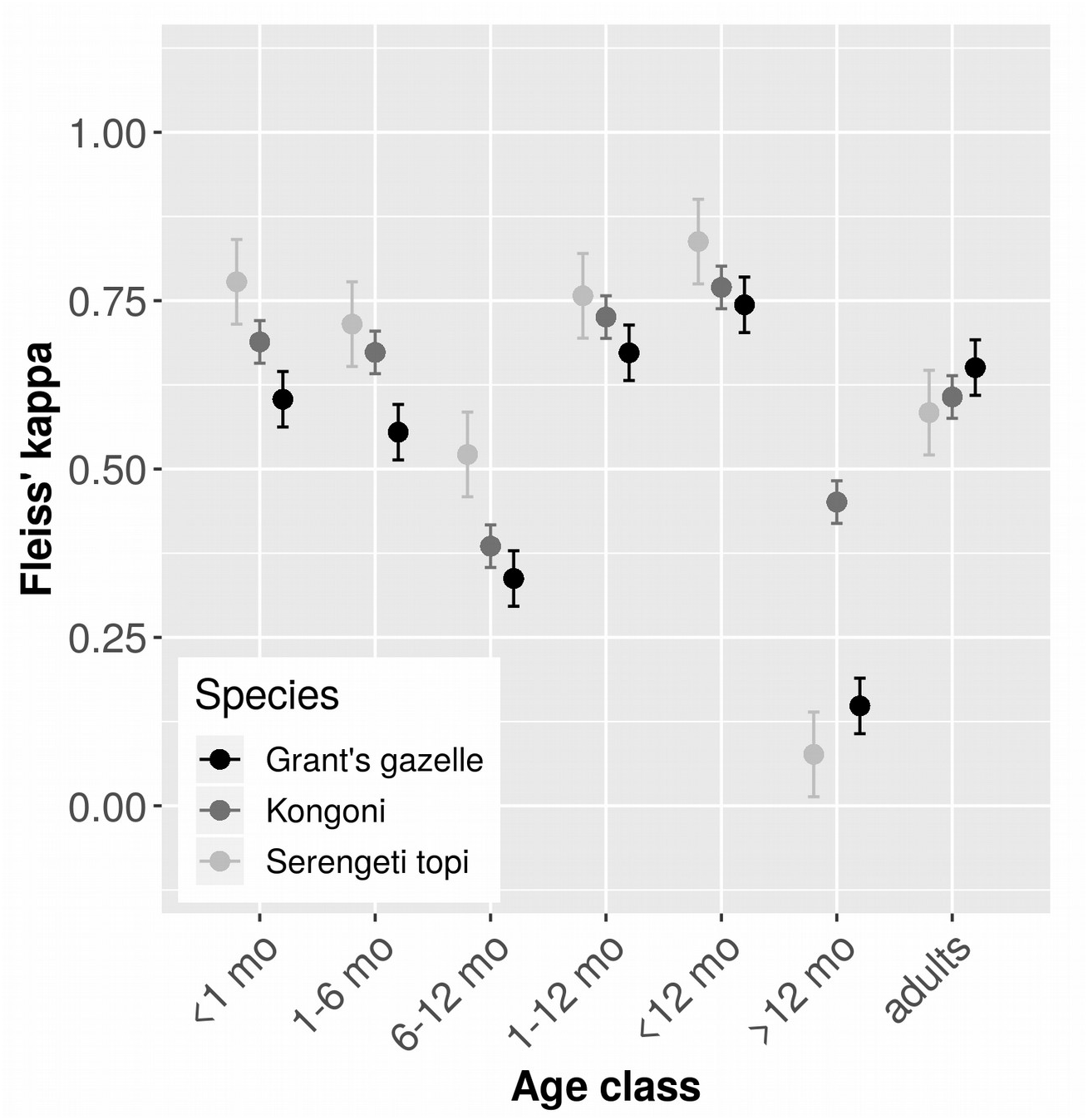
Fleiss’ κ denoting the level of agreement between the three trained observers (LT, LK and MC) in the identification of presence or absence of at least one individual belonging to any of the five age classes (< 1 month, 1-6 months, 6-12 months, > 12 months, adults), for the three species of interest. Pictures from Snapshot Serengeti Program, Tanzania, between July 2010 and April 2013. The two combined age classes “1-12 months” and “< 12 months” are also presented. Light grey dots: topi, dark grey dots: kongoni, black dots: Grant’s gazelle. Vertical bars represent the confidence intervals. A Fleiss’ κ near 1 denotes an almost perfect agreement, whereas a value near or < 0 means a very poor agreement between raters.

The model best describing our data for the age class < 1 month was the linear model for topi, the piecewise model with slopes on both sides of the threshold for kongoni and the piecewise model with slope before and plateau after the threshold for Grant’s gazelle (Table 2). The model best describing our data for the age class < 12 months was the threshold model with slopes on both sides of the threshold for topi, kongoni and Grant’s gazelle (Table 2).

**Table 2:** Statistics of models investigating the relationship between the proportion of volunteers identifying at least one “young” and the probability of presence of at least one individual < 1 or 12 months, assessed by the trained observers on a given sequence for the three species of interest: topi, kongoni and Grant’s gazelle. Best models are in bold. θ = estimated threshold (% of volunteers), QIC = Quasi-likelihood under the Independence model Criterion, in. = intercept, estimates ± standard error and [95% confidence interval].

| Age class | Species | Model type | $\theta$ | QIC | Quasi-likelihood | Estimates |
| --- | --- | --- | --- | --- | --- | --- |
| < 1 month | Topi | Null | | 924.6 | -459.77 | in. = -1.509 ± 0.133 |
|  |  | **Linear** |  | **630.9** | **-311.05** | **in. = -4.013 ± 0.342**<br>**$\beta$ = 0.048 ± 0.005** |
| | | Piecewise - slope/plateau | 9<br>0 | 635.2 | -313.23 | in. = -4.102 ± 0.362<br>$\beta_1$ = 0.051 ± 0.005<br>$\mu_2$ = 0.626 [0.523; 0.720] |
| | | Piecewise - slope/slope | 7<br>8 | 634.7 | -310.49 | in. = -8.863 ± 1.763<br>$\beta_1$ = 0.045 ± 0.007<br>$\beta_2$ = 0.064 ± 0.023 |
| | Kongoni | Null | | 2614.4 | -1304.83 | in. = -2.125 ± 0.081 |
| | | Linear | | 1993 | -992.05 | in. = -3.977 ± 0.174<br>$\beta$ = 0.043 ± 0.003 |
| | | Piecewise - slope/plateau | 9<br>0 | 2005.8 | -998.42 | in. = -4.007 ± 0.178<br>$\beta_1$ = 0.044 ± 0.003<br>$\mu_2$ = 0.498 [0.434; 0.563] |
|  |  | **Piecewise - slope/slope** | **1**<br>**0** | **1966.7** | **-977.58** | **in. = -10.522 ± 1.740**<br>**$\beta_1$ = 0.687 ± 0.178**<br>**$\beta_2$ = 0.039 ± 0.003** |
| | Grant's gazelle | Null | | 1236.2 | -615.91 | in. = -2.479 ± 0.117 |
| | | Linear | | 1016.6 | -501.81 | in. = -3.653 ± 0.181<br>$\beta$ = 0.055 ± 0.007 |
|  |  | **Piecewise - slope/plateau** | **4**<br>**3** | **969.7** | **-480.75** | **in. = -4.355 ± 0.227**<br>**$\beta_1$ = 0.097 ± 0.008**<br>**$\mu_2$ = 0.451 [0.353; 0.552]** |
| | | Piecewise -<br>slope/slope | 4<br>3 | 974.6 | -480.48 | in. = $-4.573 \pm 0.544$<br>$\beta_1 = 0.094 \pm 0.010$<br>$\beta_2 = 0.006 \pm 0.013$ |
| < 12<br>months | Topi | Null | | 1241.3 | -618 | in. = $0.698 \pm 0.111$ |
| | | Linear | | 948 | -468.64 | in. = $-0.866 \pm 0.184$<br>$\beta = 0.059 \pm 0.007$ |
| | | Piecewise -<br>slope/plateau | 6<br>0 | 964.6 | -477.23 | in. = $-0.987 \pm 0.191$<br>$\beta_1 = 0.069 \pm 0.007$<br>$\mu_2 = 0.958 [0.926; 0.977]$ |
|  |  | **Piecewise -**<br>**slope/slope** | **3**<br>**0** | **946.5** | **-464.45** | **in. = $-3.202 \pm 0.624$**<br>**$\beta_1 = 0.031 \pm 0.017$**<br>**$\beta_2 = 0.090 \pm 0.022$** |
| | Kongoni | Null | | 4992.3 | -2493.63 | in. = $0.609 \pm 0.054$ |
| | | Linear | | 3726.5 | -1858.95 | in. = $-1.139 \pm 0.108$<br>$\beta = 0.089 \pm 0.006$ |
| | | Piecewise -<br>slope/plateau | 9<br>0 | 3726.6 | -1859.03 | in. = $-1.140 \pm 0.108$<br>$\beta_1 = 0.089 \pm 0.006$<br>$\mu_2 = 0.999 [0.997; 1]$ |
|  |  | **Piecewise -**<br>**slope/slope** | **1**<br>**2** | **3684.8** | **-1835.45** | **in. = $-3.041 \pm 0.271$**<br>**$\beta_1 = 0.213 \pm 0.029$**<br>**$\beta_2 = 0.066 \pm 0.006$** |
| | Grant's<br>gazelle | Null | | 3127.4 | -1561.2 | in. = $-0.154 \pm 0.067$ |
| | | Linear | | 2940.5 | -1463.22 | in. = $-0.837 \pm 0.140$<br>$\beta = 0.049 \pm 0.010$ |
| | | Piecewise -<br>slope/plateau | 3<br>0 | 2866.2 | -1428.27 | in. = $-1.285 \pm 0.132$<br>$\beta_1 = 0.091 \pm 0.009$<br>$\mu_2 = 0.809 [0.748; 0.859]$ |
|  |  | **Piecewise -**<br>**slope/slope** | **4**<br>**1** | **2866.4** | **-1425.59** | **in. = $-0.009 \pm 0.506$**<br>**$\beta_1 = 0.083 \pm 0.010$**<br>**$\beta_2 = -0.029 \pm 0.013$** |

The probability of observing a juvenile when all volunteers reported one was near 1 for juveniles < 12 months for topi and kongoni. Concerning Grant’s gazelle, this probability only reached 0.90 when 41% of the volunteers recorded the presence of young. Between 41% and 100% of volunteers identifying young in the sequences, the probability decreased (Fig. 3 d), e), f)). When investigating the presence of juveniles < 1 month, the probability that a juvenile was actually present was never greater than 0.697 (Fig. 3 a), b), c)). On the other hand, the model predicted that when no volunteer reported the presence of juveniles, the probability that there was juveniles < 1 month was under 1.8%, but there was at least 9.6% chance to observe a juvenile < 12 months.

**Figure 3:**
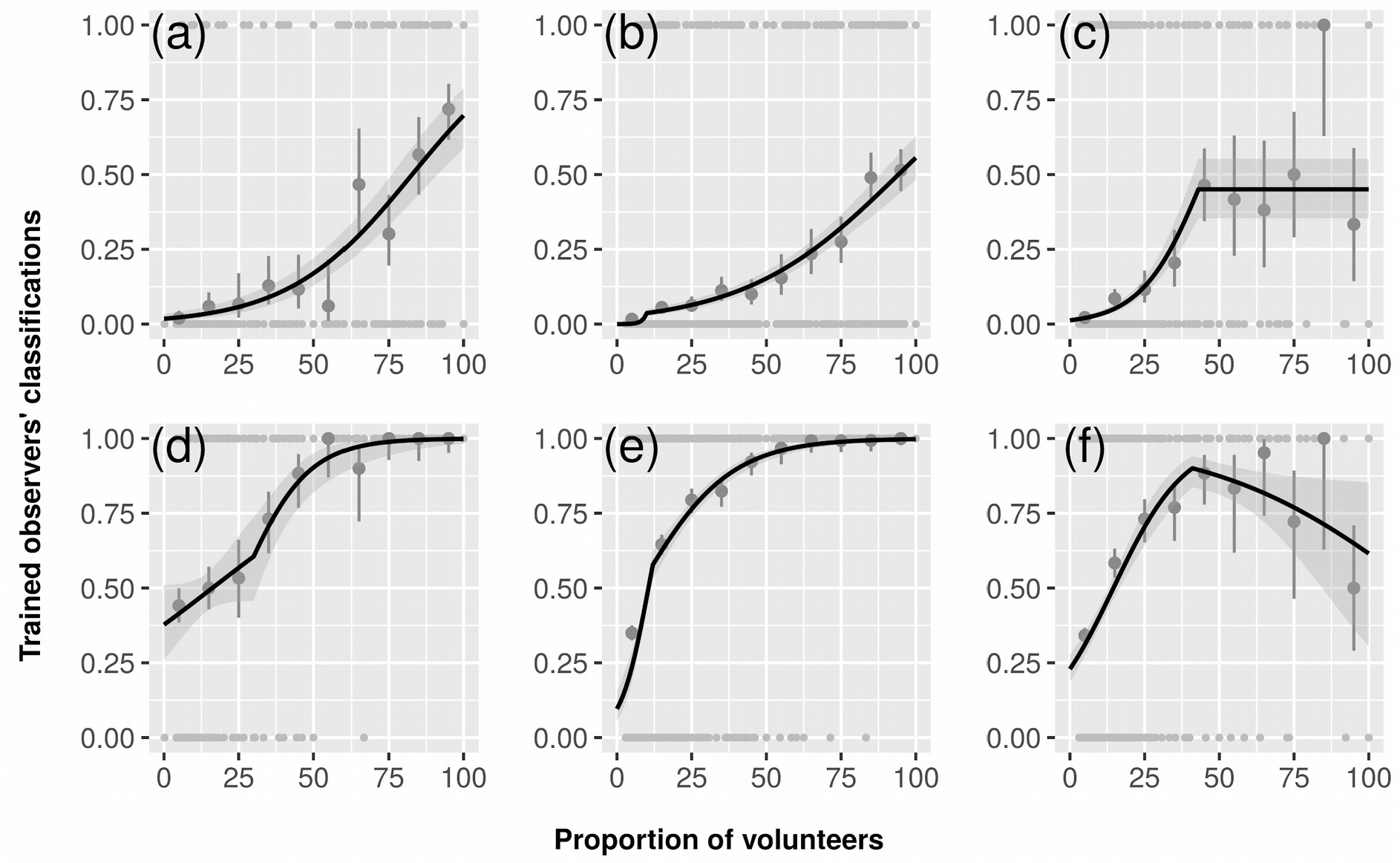
relationship, as predicted from the best model (see text for details), between the proportion of volunteers identifying the presence of “young” and the probability of presence of at least one individual < 1 (a), b), c)) and < 12 (d), e), f)) months assessed by the trained observers in a given sequence (0 < proportion of volunteers identifying “young” ≤ 1), for the three species of interest: a) and d) topi, b) and e) kongoni and c) and f) Grant’s gazelle. Pictures from Snapshot Serengeti Program, Tanzania, between July 2010 and April 2013. Light grey dots represent the probability of presence of at least one individual < selected age class in each sequence assessed by the three trained observers, dark grey dots represent the mean of those probabilities for each 10% volunteers interval, vertical bars represent 95% confidence intervals. Solid line represents predicted values from the best model. Shaded areas represent 95% confidence intervals.

Contrary to our expectations, the piecewise model characterized by two slopes was almost always selected as the best model for the three species and both age classes. This denotes a sudden change in the rate of detection from a certain percentage of volunteers. The detection rate decreases for kongoni and Grant’s gazelle for young < 12 months beyond 12 and 41% of volunteers voting for the presence of young respectively. The number of volunteers who did classify a photograph was independent from the probability to detect a young.

## Discussion

Our study reveals the strength and weaknesses of using citizen-based assessment of age classes on camera trap pictures. Clearly, citizen involvement through an online platform has been critical for the classification of the millions of photographs collected by the Snapshot Serengeti initiative. Also, despite leaving it to the volunteers to decide what a juvenile looks like, volunteers’ classification allows a rough, moderately accurate, but quick sorting of sequences with/without juveniles.

Our study makes clear that, for the species studied and given the minimal guidelines given to volunteers, the *absence* of very young (< 1 month) individuals on pictures can be reliably assumed when no volunteer reports a presence. The sequences almost never labelled by volunteers as containing “young” were very unlikely to contain young under one month of age according to the trained observers. This suggests that volunteers rarely miss very young juveniles. This likely occurs because very young juveniles are distinctively smaller than adults, and possibly also because very young mammals share some physical characteristics such as relatively large eyes, long legs, short and rounded nose, all belonging to *Kindchenschema*, known to be very attractive stimuli for humans (Brosch et al. 2007, Golle et al. 2013). By contrast, the *presence* of very young individuals appears more difficult to ascertain from volunteers’ data, and this apparently comes from the lack of guidelines given to volunteers. Indeed, consistent with the idea that volunteers easily identify very young individuals, a large consensus among volunteers around the presence of a very young juvenile could be a reliable indication of actual presence, but only when no older juveniles are present (compare Fig. 3 a), b), c) & Supporting information 4 - Fig. A). Unfortunately, the absence or presence of older juveniles cannot currently be known without a reassessment of the pictures because volunteers were not asked to differentiate between juvenile age classes. Therefore, the presence of very young juveniles remains difficult to ascertain. On the other hand, the *presence* of juveniles, irrespectively of their age, can be reliably assumed when all volunteers agree about a presence (Fig. 3 a), b), c)), especially for topi and kongoni: the model predicted that when all the volunteers reported the presence of juveniles, the probability that there was juveniles < 12 months was indeed at least 99.8%. This time, the *absence* of juveniles regardless of their age is less accurate, meaning that they are missed quite often. As they grow, juveniles become increasingly similar to adults, and are more likely to be mistaken for the latter by volunteers. Anyhow, we emphasize that detectability of juveniles is not equal between species (Fig. 3).

Among the three trained observers, we found the best agreement in the detection of age classes for a given sequence for topi and kongoni, suggesting that they are the easiest species to classify, and Grant’s gazelle the hardest. The combination of small body size and gregariousness could explain why the determination of age was more difficult and heterogeneous for Grant’s gazelle than for the two other species. Topi and kongoni are fairly large, and small body size makes the detection of some inconspicuous age criteria challenging (*e.g.* presence or absence of very small horn buds on the forehead of the individuals). On the contrary, it is certainly easier to discriminate between adults and juveniles in the largest herbivores, such as giraffe (*Giraffa camelopardalis*) or elephant (*Loxodonta africana*) because of the marked size difference between them. Discrimination between several juvenile age classes will certainly be difficult still as individuals are often only partially spotted by camera traps and subtle aging criteria are not visible. More generally, in species forming large herds such as wildebeest, young might be particularly difficult to spot and are likely missed frequently. In large groups, the body of many individuals overlaps, hampering our ability to see aging criteria correctly, and to age them accurately. This could explain why in our study for instance, Grant’s gazelle frequently occurring in large herds reaching more than 30 individuals in some sequences, is also the one with the lowest identification success of juveniles. Finally, a pronounced sexual dimorphism in horn growth in Grant’s gazelle likely led to confusions between the first two age classes as male horns grow faster, a young male can look like an older female when using the length of the horns as an aging criteria. This would also be the case for species with a sexual dimorphism in which only males grow horns like impala (*Aepyceros melampus*) or waterbuck (*Kobus defassa*). Clearly, a description of the morphological changes that occur throughout the development of young herbivores (*e.g.* Spinage 1976 in our species, Cunningham et al. 2011, Dezeure et al. 2020 in other species) is of great value and substantially helps at reaching consistent results among different observers.

Another limitation of camera trapping in the context of reproductive studies is the level of effort required to obtain a sufficient sample size. Here, working with data from one of the world’s largest and long-running camera trap studies, we ended up with a small number of sequences with juveniles < 1 month old after appropriate data selection. We identified four main explanations for this important reduction in exploitable sequences. First and foremost, our ability to determine an individual’s age class depends strongly on the photograph quality and particularly its framing and exposure. In many cases, individuals were located too far from the camera or were only partly visible, or photographs were too blurry, dark, or overexposed to be scrutinized, leading to potentially significant loss of reproductive data and a high number of individuals of unknown or over-estimated age. Another potential source of information loss was species misidentification by volunteers. In our study, about 30% of the sequences labelled with Grant’s gazelle were misidentified because it greatly resembles species such as impala and Thomson’s gazelle which are also present in the study site. Species abundance obviously directly impacts the number of sequences collected. The abundance of the three studied species is low compared to other ungulates in the Serengeti system. Sinclair reported 55,500 individuals topi, 20,700 individuals kongoni and 6,000 individuals Grant’s gazelle in the 1970s, whereas the numbers of wildebeest and zebra were 720,000 and 240,000 individuals respectively (Sinclair and Norton-Griffiths 1995). Finally, the anti-predator strategy of juvenile large herbivores, known as the hider-follower gradient (Lent 1974, Rutberg 1987), could influence the number of sequences containing very young individuals. While followers become active and stick with their mother just a few hours after birth, hiders stay concealed in dense vegetation during their first weeks of life. The detection probability of hiders from camera traps should then be much lower than of followers, consistent with our observation of very young topi seen in a greater proportion than kongoni and Grant’s gazelle.

Volunteer classification provides information that can reliably be used to infer the *presence* of juveniles < 12 months or the *absence* of juveniles < 1 month, as these annotations appear robust. However, because volunteers seem able to discriminate between individuals of less than one month old and the rest of the juveniles even in the absence of any stated criteria, more precise results could be achieved by asking them to differentiate between two age classes of juveniles, such as “juvenile” and “newborn”. The level of agreement between trained observers in the classification of age classes according to species is also a good indication of what kind of tasks could be successfully conducted by volunteers. When this agreement is low (*e.g.* in the identification of topi and Grant’s gazelle yearlings), one could not expect volunteers to properly identify such an age class. We advise to limit the number of classes the volunteers are asked to identify, and to focus on the most recognisable ones. Another way to improve results generated via citizen science platforms could be the inclusion of detailed information, as for species identification (Swanson et al. 2015). When a volunteer detects a young on his/her photograph, he or she could be prompted with comprehensive keys to age juvenile from its morphology along with a set of reference pictures or drawings.In general, however, we would recommend to ask citizen scientists to identify newborns, *i.e.* individuals under one month of age vs. other juveniles. To identify newborns of bovid species with horns like in our study, volunteers would have to look for the smallest individuals (the head does not come above the back of the female, Ogutu et al. 2008), with no evidence of bud horns, with specific coat color (*e.g.* darker coat color in Grant’s gazelle in our study, or lighter coat color in wildebeest) or even with umbilical cord remnants. One difficulty is to adapt the different criteria to the every species studied.

We finally suggest evaluating volunteers’ classification skills by presenting them with images of individuals of known age (captive or tagged animals for instance) and assessing their accuracy compared to labels provided by experts. Volunteers could then be assigned classification tasks adapted to their skills (*e.g.* species identification would belong to the easiest tasks, whereas age classification would belong to the hardest). Snapshot Safari and Zooniverse continue to create new modes of annotation that best leverage the public’s interest in contributing to research, and this is a logical next step for the Snapshot Safari initiative. Overall, we find that by closely investigating the data already collected by volunteer-based programs, data collection procedures can be adjusted to enhance the contributions of citizen scientists to scientific research and conservation efforts.

## Declarations

## Acknowledgments

A PhD fellowship from the university Lyon 1 attributed to LT. This work was performed using the computing facilities of the CC LBBE/PRABI. The authors thank T. Michael Anderson for maintaining the camera traps in Tanzania, Zooniverse for hosting “Snapshot Serengeti”, >130,000 online volunteers for classifying the images, the Tanzania Wildlife Research Institute and Tanzania National Parks for research permission, and the Minnesota Supercomputing Institute (http://www.msi.umn.edu) for contributing to data storage/processing and analysis.

## Authors’ contributions

LT, CB and SCJ conceived the ideas and designed methodology; CP and SH organized data collection; LT, LK and MC analysed the data; LT, CB and SCJ led the writing of the manuscript. All authors contributed critically to the drafts and gave final approval for publication.

## Data availability

Pictures available from the Labeled Image Library of Alexandria – Biology and Conservation: http://lila.science/datasets/snapshot-serengeti and data from the Dryad Data Repository: *to be completed*.

