## Supplementary material for "Can citizen science analysis of camera trap data be used to study reproduction? Lessons from Snapshot Serengeti program"

### Supporting information

Appendix 1: number of sequences per species and per year, according to the classifiers: zooniverse volunteers and trained observers as LT, LK and MC (pictures from Snapshot Serengeti program, Tanzania, between July 2010 and April 2013). Here is reported the number of sequences only for sequences gathering at least one volunteer identifying at least one juvenile on the sequence. Note that years go from July of the year *n* to June of the following year *n+1*.

| **Species** | **Year** | **Total number of sequences assessed by the volunteers** | **Total number of sequences assessed by the trained observers after reclassification of the right species** |
|---|---|---|---|
| **Topi** | Total | 281 | 324 |
| | July 2010 - June 2011 | 101 | 105 |
| | July 2011 - June 2012 | 78 | 98 |
| | July 2012 - June 2013 | 102 | 121 |
| **Kongoni** | Total | 1,290 | 1,281 |
| | July 2010 - June 2011 | 412 | 433 |
| | July 2011 - June 2012 | 501 | 489 |
| | July 2012 - June 2013 | 377 | 359 |
| **Grant's gazelle** | Total | 1,095 | 754 |
| | July 2010 - June 2011 | 491 | 358 |
| | July 2011 - June 2012 | 368 | 221 |
| | July 2012 - June 2013 | 236 | 175 |

Appendix 2: Selection criteria to decide on the studied species (see text for details). Pictures from Serengeti Snapshot Program, Tanzania, between July 2010 and April 2013. "+" corresponds to a satisfied criterion, "-" corresponds to an unsatisfied criterion, the choice is based on informal criteria. The species studied here, which fulfilled the most criteria, are in bold.

| **Species** | **Nb of sequences available** | **Presence of horns in males and females** | **Large size of the young** | **Small group size** |
|---|---|---|---|---|
| African Buffalo | + | + | + | - |
| Blue Wildebeest | - | + | + | - |
| Bushbuck | - | - | - | + |
| Dik-dik | + | - | - | + |
| Eland | - | + | + | + |
| Elephant | + | - (but tusks) | + | + |
| Giraffe | + | - | + | + |
| **Grant's gazelle** | **+** | **+** | **+** | **+** |
| Hippopotamus | - | - | + | + |
| Impala | + | - | + | - |
| **Kongoni** | **+** | **+** | **+** | **+** |
| Reedbuck | + | - | + | + |
| Rhinoceros | - | + | + | + |
| Thomson's gazelle | - | + | - | - |
| **Topi** | **+** | **+** | **+** | **+** |
| Warthog | + | - (but tusks) | - | + |
| Waterbuck | - | - | + | + |
| Zebra | - | - | + | - |

Appendix 3: Criteria for age classification of topi, kongoni and Grant's gazelle.

| **Species** | **Category** | **Criterion** | **Age** | **Reference** |
|---|---|---|---|---|
| Topi | *Horns* | Absence of horns | newborn | Ogutu et al. 2010 |
| | | Horns visible but shorter than the ear | less than 6 months | Ogutu et al. 2008 |
| | | Presence of "bud" horns | 6 months | Jewell 1972 |
| | | Horns still straight; less than twice the length of the ear | less than 10 months | Ogutu et al. 2008 |
| | | 7 to 9 ridges on the horns | more than 12 months | Jewell 1972 |
| | *Coat color* | Pale fawn coat | 6 months | Jewell 1972 |
| | *Body size* | Half adult size | 6 months | Jewell 1972 |
| Kongoni | *Horns* | Absence of horns | newborn to 1 month | Gosling 1969 |
| | | Horns straight, approximately 10 cm | 2 or 3 months | Ansell 1960 |
| | | Horns visible but shorter than the ear | less than 6 months | Ogutu et al. 2008 |
| | | Horns straight, less than twice the length of the ear | less than 10 months | Ogutu et al. 2008 |
| | | Horns distinctly curved in toward each other | 11 to 18 months | Ogutu et al. 2008 |
| | | Horns lyre-shape but not full size | 19 to 24 months | Ogutu et al. 2008 |
| Grant's gazelle | *Horns* | In males and females: top of the horns visible | 5 to 6 months | Walther 1972 |
| | | In males: horns are ⅓ of ear length, clearly more massive than adult female's one, forehead at the basis of the horns swollen (corresponding to the | 7 to 12 months | Walther 1972 |

| | | | | |
|---|---|---|---|---|
| | | eruption of the first ring(s)) | | |
| | | In males: horns as long as in adult females or longer, with pronounced rings, but thicker than in adult females. No backward curvature yet, but rather concavely and a little inward curved at the top | 12 to 24 months | Walther 1972 |
| | | In females: length of the horns slightly below those of an adolescent male, and much thinner | 7 to 12 months | Walther 1972 |
| | | In females: horns longer than the ear but by far not as long as in a fully adult female | less than 24 months | Walther 1972 |
| | *Coat color* | darker in color | newborn to 2 weeks | Walther 1972 |
| | *Body size* | In males and females: half the size of an adult female, or a little more | 5 to 6 months | Walther 1972 |
| | | In males: size about ¾ of that of adult male, body still weaker than adult female's one | 7 to 12 months | Walther 1972 |
| | | In males: size and body strength of an adult female | 12 to 24 months | Walther 1972 |
| | | In females: approximately, although not yet completely, reached the size of an adult female, body still slimmer | 12 to 24 months | Walther 1972 |

| | | | | |
|---|---|---|---|---|
| | *General* | 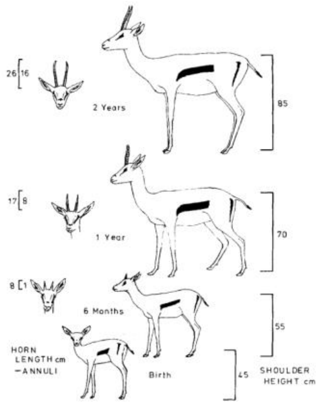 | | Spinage 1976 |
| | *General* | 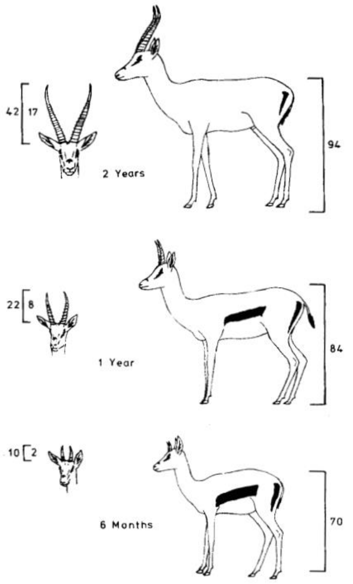 | | Spinage 1976 |

For all the species: persistence of the umbilical cord denotes very recent birth (*e.g.* Ogutu et al. 2010).

Appendix 4: Evaluation of the significance of the difference between the Fleiss' κ values by the method of the bootstraps.

To check for the significance of the difference between the Fleiss' κ values between species for a given age class and between ages classes for a given species, we used the bootstraps methods described in Vanbelle and Albert (2008). We describe here the estimation of the difference between species for a given age class, but the procedure is the same for the estimation of the difference between ages classes for a given species.

We generated 9,999 batch (the $10{,}000^{ths}$ corresponding to the Fleiss' κ calculated from the original dataset) by sampling with replacement n=324, 1,281 and 754 sequences for the topi, the kongoni and he Grant's gazelle respectively, corresponding to the number of sequences with at least one individual of known age. We calculated the difference in Fleiss' κ values between species, two by two for a given age class. We then applied a one sample Student's test on the distribution of the 10,000 differences with α level = 5% to check if it is significantly different from 0.

The results of the comparison of the Fleiss' κ values between species for a given age class are reported here (significant p-values are in bold type):

| **Species compared** | **Age class** | **p value** | **CI lower boundary** | **CI upper boundary** | **t statistic** |
|---|---|---|---|---|---|
| Topi - Kongoni | < 1 month | < **0.001** | 0.08754 | 0.08936 | 189.89 |
| | 1 - 6 months | < **0.001** | 0.04046 | 0.04183 | 117.28 |
| | 6 - 12 months | < **0.001** | 0.13225 | 0.135 | 190.24 |
| | 1 - 12 months | < **0.001** | 0.03029 | 0.03156 | 94.89 |
| | < 12 months | < **0.001** | 0.06714 | 0.06832 | 226.57 |

| | | | | | |
|---|---|---|---|---|---|
| | > 12 months | **< 0.001** | -0.37617 | -0.37418 | -737.73 |
| | adults | **< 0.001** | -0.02906 | -0.02641 | -41.06 |
| Topi - Grant's gazelle | < 1 month | **< 0.001** | 0.1739 | 0.17617 | 302.23 |
| | 1 - 6 months | **< 0.001** | 0.15953 | 0.16111 | 397.07 |
| | 6 - 12 months | **< 0.001** | 0.18003 | 0.18283 | 253.83 |
| | 1 - 12 months | **< 0.001** | 0.08313 | 0.08453 | 234.12 |
| | < 12 months | **< 0.001** | 0.09347 | 0.09473 | 292.15 |
| | > 12 months | **< 0.001** | -0.07374 | -0.0719 | -155.15 |
| | adults | **< 0.001** | -0.07205 | -0.0693 | -100.8 |
| Kongoni - Grant's gazelle | < 1 month | **< 0.001** | 0.08557 | 0.08759 | 168.13 |
| | 1 - 6 months | **< 0.001** | 0.11858 | 0.11977 | 393.43 |
| | 6 - 12 months | **< 0.001** | 0.04689 | 0.04871 | 103.03 |
| | 1 - 12 months | **< 0.001** | 0.05239 | 0.05342 | 199.75 |
| | < 12 months | **< 0.001** | 0.02589 | 0.02685 | 108.39 |
| | > 12 months | **< 0.001** | 0.30152 | 0.3032 | 706.03 |
| | adults | **< 0.001** | -0.04361 | -0.04227 | -125.74 |

The results of the comparison of the Fleiss' κ values between age classes for a given species are reported here (significant p-values are in bold type):

| **Age classes compared** | **Species** | **p value** | **CI lower boundary** | **CI upper boundary** | **t statistic** |
|---|---|---|---|---|---|

| | | | | | |
|---|---|---|---|---|---|
| < 1 month / 1-6 months | Topi | < **0.001** | 0.06109 | 0.06301 | 127.16 |
| | Kongoni | < **0.001** | 0.01413 | 0.01536 | 46.93 |
| | Grant's gazelle | < **0.001** | 0.04635 | 0.04833 | 93.74 |
| < 1 month / 6-12 months | Topi | < **0.001** | 0.25694 | 0.2598 | 354.08 |
| | Kongoni | < **0.001** | 0.30272 | 0.30437 | 721.38 |
| | Grant's gazelle | < **0.001** | 0.26369 | 0.26583 | 484.89 |
| < 1 month / 1-12 months | Topi | < **0.001** | 0.01931 | 0.02117 | 42.63 |
| | Kongoni | < **0.001** | -0.0379 | -0.03667 | -118.61 |
| | Grant's gazelle | < **0.001** | -0.07192 | -0.07 | -144.77 |
| < 1 month / < 12 months | Topi | < **0.001** | -0.06178 | -0.05999 | -133.34 |
| | Kongoni | < **0.001** | -0.08221 | -0.08099 | -262.32 |
| | Grant's gazelle | < **0.001** | -0.14276 | -0.14087 | -294.92 |
| < 1 month / > 12 months | Topi | < **0.001** | 0.70058 | 0.70267 | 1317.59 |
| | Kongoni | < **0.001** | 0.23715 | 0.23886 | 546.53 |
| | Grant's gazelle | < **0.001** | 0.45276 | 0.45479 | 877.02 |
| < 1 month / adults | Topi | < **0.001** | 0.19596 | 0.19887 | 265.28 |
| | Kongoni | < **0.001** | 0.08056 | 0.08189 | 238.15 |
| | Grant's gazelle | < **0.001** | -0.04931 | -0.04728 | -93.38 |
| 1-6 months / 6-12 months | Topi | < **0.001** | 0.19494 | 0.1977 | 279 |

| | | | | | |
|---|---|---|---|---|---|
| | Kongoni | < **0.001** | 0.28809 | 0.2895 | 802.83 |
| | Grant's gazelle | < **0.001** | 0.2166 | 0.21825 | 515.13 |
| 1-6 months / 1-12 months | Topi | < **0.001** | -0.04264 | -0.04098 | -98.36 |
| | Kongoni | < **0.001** | -0.05246 | -0.0516 | -235.71 |
| | Grant's gazelle | < **0.001** | -0.11895 | -0.11764 | -354.23 |
| 1-6 months / < 12 months | Topi | < **0.001** | -0.12373 | -0.12214 | -303.89 |
| | Kongoni | < **0.001** | -0.09678 | -0.09592 | -440.62 |
| | Grant's gazelle | < **0.001** | -0.18978 | -0.18852 | -589.71 |
| 1-6 months / > 12 months | Topi | < **0.001** | 0.63861 | 0.64055 | 1291.37 |
| | Kongoni | < **0.001** | 0.22254 | 0.22398 | 607.57 |
| | Grant's gazelle | < **0.001** | 0.40572 | 0.40716 | 1102.75 |
| 1-6 months / adults | Topi | < **0.001** | 0.13396 | 0.13677 | 188.26 |
| | Kongoni | < **0.001** | 0.06596 | 0.067 | 251.53 |
| | Grant's gazelle | < **0.001** | -0.09636 | -0.0949 | -257.31 |
| 6-12 months / 1-12 months | Topi | < **0.001** | -0.23947 | -0.23678 | -346.8 |
| | Kongoni | < **0.001** | -0.34152 | -0.34014 | -968.91 |
| | Grant's gazelle | < **0.001** | -0.33651 | -0.33494 | -840.16 |
| 6-12 months / < 12 months | Topi | < **0.001** | -0.32057 | -0.31793 | -475.15 |
| | Kongoni | < **0.001** | -0.38584 | -0.38446 | -1094.11 |

| | | | | | |
|---|---|---|---|---|---|
| | Grant's gazelle | < **0.001** | -0.40734 | -0.40582 | -1047.05 |
| 6-12 months / > 12 months | Topi | < **0.001** | 0.44182 | 0.44469 | 605.49 |
| | Kongoni | < **0.001** | -0.06645 | -0.06463 | -141.54 |
| | Grant's gazelle | < **0.001** | 0.18816 | 0.18987 | 435.7 |
| 6-12 months / adults | Topi | < **0.001** | -0.06273 | -0.05917 | -67.15 |
| | Kongoni | < **0.001** | -0.22307 | -0.22156 | -577.94 |
| | Grant's gazelle | < **0.001** | -0.3139 | -0.31221 | -728.07 |
| 1-12 months / < 12 months | Topi | < **0.001** | -0.08188 | -0.08037 | -210.52 |
| | Kongoni | < **0.001** | -0.04473 | -0.0439 | -208.96 |
| | Grant's gazelle | < **0.001** | -0.07142 | -0.07028 | -244.31 |
| 1-12 months / > 12 months | Topi | < **0.001** | 0.68045 | 0.68232 | 1425.68 |
| | Kongoni | < **0.001** | 0.27457 | 0.27601 | 747.8 |
| | Grant's gazelle | < **0.001** | 0.52407 | 0.52541 | 1531.1 |
| 1-12 months / adults | Topi | < **0.001** | 0.17579 | 0.17857 | 249.87 |
| | Kongoni | < **0.001** | 0.11801 | 0.11902 | 459.96 |
| | Grant's gazelle | < **0.001** | 0.022 | 0.02334 | 66.05 |
| < 12 months / > 12 months | Topi | < **0.001** | 0.76161 | 0.76341 | 1653.02 |
| | Kongoni | < **0.001** | 0.31889 | 0.32032 | 873.98 |
| | Grant's gazelle | < **0.001** | 0.59494 | 0.59624 | 1792.69 |

| | | | | | |
|---|---|---|---|---|---|
| < 12 months / adults | Topi | **< 0.001** | 0.25694 | 0.25966 | 371.9 |
| | Kongoni | **< 0.001** | 0.16233 | 0.16333 | 641.84 |
| | Grant's gazelle | **< 0.001** | 0.09287 | 0.09417 | 281 |
| > 12 months / adults | Topi | **< 0.001** | -0.50568 | -0.50275 | -673.85 |
| | Kongoni | **< 0.001** | -0.15755 | -0.15601 | -399.09 |
| | Grant's gazelle | **< 0.001** | -0.50281 | -0.50133 | -1322.38 |

Appendix 5: Detection of juveniles < 1 month when no other juveniles are present.

Concerning juveniles < 1 month, we developed two kinds of models: one with the complete dataset (see "Material and methods" and "Results"), and another one exclusively with sequences characterized by 1) a perfect agreement between the three trained observers concerning the presence or absence of the juveniles < 1 month, 2) a perfect agreement between the three trained observers concerning the absence of other categories of juveniles. This second model was developed to make sure that the other juvenile age classes did not interfere with our definition of "no juvenile in the sequence" as it is not necessarily the same for trained observers (it means the absence of juveniles < 1 month old) and for the volunteers (it means the absence of "young"). So we could analyse both the extent to which volunteers classification can be used to infer on the presence of juveniles < 1 month on the one hand (model on the complete dataset presented in "Results"), and the ability of the volunteers to detect juveniles < 1 month on the other hand (model on selected dataset presented here, sequences containing juveniles older than one month removed). We fitted the same models as presented in "Material and methods".

In the absence of any other juvenile, volunteers are able to detect very precisely the presence or absence of juveniles < 1 month in topi and kongoni (Fig. A). This ability is less clear for Grant's gazelle, as the probability to effectively observe juveniles < 1 month when all the volunteers agree about the presence of juveniles is only 0.670. It seems reasonable to think that one could achieve precise estimation of the presence of very young individuals by asking volunteers to identify two juvenile age classes, such as "juvenile" and "newborn".

Table A: statistics of models investigating the relationship between the proportion of volunteers identifying at least one "young" and the probability of presence of at least one individual < 1 month, assessed by the trained observers on a given sequence for the three species of interest: topi, kongoni and Grant's gazelle (on a selected dataset, see text for details). Best models are in bold. $\theta$ = estimated threshold (% of volunteers), *QIC* = Quasi-likelihood under the Independence model Criterion, *in.* = intercept, estimates ± standard error and [95% confidence interval].

| **Age class** | **Species** | **Model type** | **θ** | **QIC** | **Quasi-likelihood** | **Estimates** |
|---|---|---|---|---|---|---|
| < 1 month gold | Topi | Null | | 163.2 | -80.61 | *in.* = -0.822 ± 0.190 |
| | | **Linear** | | **44.1** | **-18.43** | ***in.* = -5.378 ± 0.816**<br>**□ = 0.116 ± 0.022** |
| | | Piecewise - slope/plateau | 80 | 44 | -18.78 | *in.* = -5.589 ± 0.844<br>□*1* = 0.124 ± 0.020<br>$\mu2$ = 0.987 [0.907; 0.998] |
| | | Piecewise - slope/slope | 22 | 44.2 | -17.8 | *in.* = -10.768 ± 1.585<br>□*1* = 0.288 ± 0.067<br>□*2* = 0.097 ± 0.021 |
| | Kongoni | Null | | 376.5 | -187.26 | *in.* = -1.621 ± 0.132 |
| | | Linear | | 100.6 | -47.64 | *in.* = -5.370 ± 0.474<br>□ = 0.139 ± 0.016 |
| | | Piecewise - slope/plateau | 77 | 100.6 | -47.81 | *in.* = -5.440 ± 0.467<br>□*1* = 0.142 ± 0.014<br>$\mu2$ = 0.996 [0.980; 0.999] |
| | | **Piecewise - slope/slope** | **14** | **98.1** | **-45.71** | ***in.* = -10.042 ± 1.889**<br>**□*1* = 0.415 ± 0.149**<br>**□*2* = 0.108 ± 0.015** |
| | Grant's gazelle | Null | | 166.8 | -82.42 | *in.* = -2.714 ± 0.220 |
| | | Linear | | 116.2 | -52.88 | *in.* = -4.271 ± 0.389<br>□ = 0.071 ± 0.018 |
| | | **Piecewise -** | **39** | **88.3** | **-41.58** | ***in.* = -6.032 ± 0.669** |

| | | | | | | |
|---|---|---|---|---|---|---|
| | | **slope/plateau** | | | | **□*1* = 0.173 ± 0.022**<br>***μ2* = 0.670 [0.480; 0.816]** |
| | | Piecewise - slope/slope | 39 | 90.2 | -41.33 | *in.* = -5.695 ± 0.852<br>□*1* = 0.185 ± 0.027<br>□*2* = -0.013 ± 0.017 |

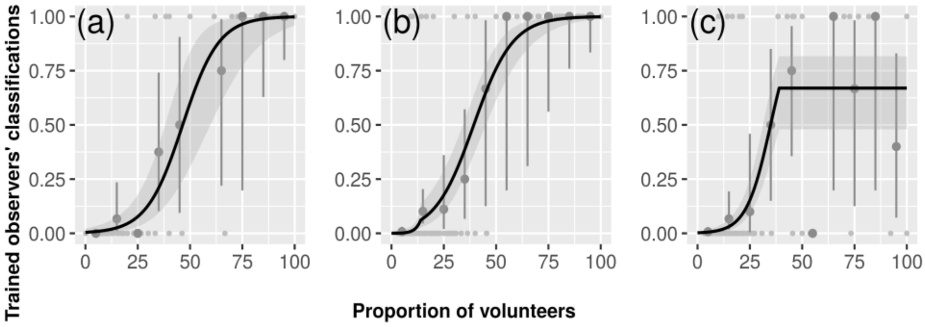


Figure A: relationship, as predicted from the best model (see text for details), between the proportion of volunteers identifying the presence of "young" and the probability of presence of at least one individual < 1 month assessed by the trained observers in a given sequence on a selected dataset (0 < proportion of volunteers identifying "young" ≤ 1, sequences containing juveniles older than one month removed), for the three species of interest: a) topi, b) kongoni and c) Grant's gazelle. Light grey dots represent the probability of presence of at least one individual < 1 month in each sequence assessed by the three trained observers, dark grey dots represent the mean of those probabilities for each 10% volunteers interval, vertical bars represent 95% confidence intervals. Solid line represents predicted values from the best model. Shaded areas represent 95% confidence intervals.

References appendices:


Ansell, W. 1960. The breeding of some larger mammals in northern rhodesia. – Proc. Zool. Soc. Lond. 134: 251–274. https://doi.org/10.1111/j.1469-7998.1960.tb05592.x

Gosling, L. 1969. Parturition and related behaviour in coke's hartebeest, alcelaphus buselaphus cokei günther. – J. Reprod. Fertil. Suppl. 6: 265–286

Jewell, P. 1972. Social organisation and movements of topi (damaliscus korrigum) during the rut, at ishasha, queen elizabeth park, uganda. – Afr. Zool. 7(1): 233–255. https://doi.org/10.1080/00445096.1972.11447442

Ogutu, J. O. et al. 2008. Rainfall influences on ungulate population abundance in the mara-serengeti ecosystem. – J. Anim. Ecol. 77(4): 814–829. https://doi.org/10.1111/j.1365-2656.2008.01392.x

Ogutu, J. O. et al. 2010. Rainfall extremes explain interannual shifts in timing and synchrony of calving in topi and warthog. – Popul. Ecol. 52(1): 89–102. https://doi.org/10.1007/s10144-009-0163-3

Spinage, C. 1976. Age determination of the female grant's gazelle. – Afr. J. Ecol. 14(2): 121–134. https://doi.org/10.1111/j.1365-2028.1976.tb00157.x

Vanbelle, S. and Albert, A. 2008. A bootstrap method for comparing correlated kappa coefficients. – J. Stat. Comput. Simul. 78(11): 1009–1015. https://doi.org/10.1080/00949650701410249

Walther, F. R. 1972. Social grouping in Grant's gazelle (gazella granti brooke 1827) in the serengeti national park. – Z. Tierpsychol. 31(4): 348–403. https://doi.org/10.1111/j.1439-0310.1972.tb01775.x
